# CircDiscoverer: A multispecies comprehensive resource for circRNA–protein interactions and RNA modification landscapes

**DOI:** 10.64898/2026.01.08.698416

**Authors:** Sreeram Srinivasan, Shivam Kumar, Samrat Chatterjee, Ajit Chande

## Abstract

Circular RNAs (circRNAs) regulate various cellular processes by interacting with miRNAs, proteins, and by encoding peptides. Compared to circRNA-miRNA interactions, circRNA-protein interactions (CPIs) and circRNA modifications have been relatively less explored. Here, we developed a comprehensive user-friendly web resource, CircDiscoverer, to explore CPIs and modification landscapes across multiple species, including *Homo sapiens, Mus musculus, Drosophila melanogaster, and Arabidopsis thaliana*. CircDiscoverer integrates manually curated literature-supported CPIs with computationally predicted interactions. Furthermore, it offers comprehensive insights into protein-binding profiles and detailed information on interacting circRNAs, including qRT-PCR primer sequences, guide RNAs for experimental validation and potential RNA modifications, thereby providing an integrated platform for further research. CircDiscoverer is available at https://aclab.iiserb.ac.in.

## 1. Introduction

Circular RNAs (circRNAs) have emerged as an important class of regulatory transcripts involved in gene regulation, diseases, and physiological processes by interacting with miRNAs and proteins as well as by encoding proteins(1–5). Although early circRNA research primarily focused on miRNA sponging(6–8), increasing evidence suggests that circRNA–protein interactions (CPIs) form stable complexes that are resistant to exonuclease degradation and influence downstream cellular functions(9, 10). While most CPI studies have been reported in humans, emerging evidence from model organisms, such as *Mus musculus* and *Drosophila melanogaster*, suggests that CPIs may represent conserved regulatory mechanisms across species(11, 12). These organisms also provide important experimental systems to investigate circRNA biology and functional CPI networks. However, the large-scale experimental validation of CPIs is labor-intensive, time-consuming, and resource-demanding, highlighting the need for integrative computational resources for systematic CPI exploration.

Although several circRNA resources exist (Table 1), CPI-centered databases remain limited and fragmented. Existing resources for CPI currently provide limited coverage and support for experimental validation, protein-centric exploration, or contextual annotation of candidate circRNAs (Table 2). This creates a bottleneck for both computational and experimental researchers attempting to prioritize CPI candidates and translate in silico predictions into bench-ready validation strategies.

To address this gap, we developed CircDiscoverer, a multispecies CPI resource integrating literature-supported and computationally predicted interactions across *Homo sapiens*, *Mus musculus*, *Drosophila melanogaster*, and *Arabidopsis thaliana*. To support the validation of the predicted CPIs, CircDiscoverer offers qRT-PCR primers, gRNAs, predicted and experimentally validated RNA modification annotations, protein-binding regions, and RNA-binding motifs, that is, short-sequence elements recognized by RNA-binding proteins (RBPs) within candidate circRNAs. Together, CircDiscoverer enables the functional exploration of circRNAs and facilitates the transition from computational CPI prediction to experimental validation.

## 2. Materials and methods

### CPI data integration strategy

CPIs were predicted using two different strategies. First, publicly available RIP-seq and CLIP-seq datasets of all the proteins from NCBI-SRA were analysed using CIRI2 and circExplorer2(13, 14) to identify circRNAs by aligning to the respective genome (Additional File 1), followed by retaining the circRNAs present in circAtlas 3.0(15) (For Homo sapiens and Mus musculus), CIRCPedia 3.0(16) (For Drosophila Melanogaster), and PlantCircRNA (17) (For Arabidopsis thaliana). Second, for proteins with experimentally characterized RNA-binding motifs(18), motif instances were scanned across circRNAs detected using CLIP-seq and RIP-Seq to provide an additional motif-supported line of evidence.

### Literature Collection

Literatures of CPIs were gathered from <u>PubMed</u> using the keywords “**protein**” and “**circular RNA**.” The retrieved articles were manually inspected for accuracy. The curated CPI information was then incorporated into the CircDiscoverer.

### Experimental design annotations

To facilitate downstream validation of the CPIs, Back-Splice-junction (BSJ)-centered qRT-PCR primers and gRNAs were designed. Briefly, we used our own custom previously designed <u>circDR-Seq script</u>(19) to generate shortened backsplice junction sequences of the mature circRNAs, and then designed BSJ-spanning qRT-PCR primers using a command line-based primer3(20), followed by curating additional experimentally validated primers from the CircAD database(21). gRNAs targeting BSJs were designed for RfxCas13d-based perturbation using the criteria described in the manuscript(22), and possible RNA modifications were inferred by overlapping circRNA coordinates with modification annotations from RMBase v.3.0(23), followed by sourcing experimentally supported records from m6A2Target(24) and other literature.

### Web implementation

CircDiscoverer was developed using Dash, which is a productive Python framework for building interactive web applications. The front-end was customized using HTML, CSS, and JavaScript to provide a responsive and user-friendly design. Flask and JavaScript were used in the backend to manage server-side logic and data flow. All data were stored and retrieved using the MongoDB database, enabling the efficient handling of genomic datasets. The platform was designed to ensure smooth interaction, high performance, and intuitive data visualization for user needs.

## 3. Results

### Resource scope and content

CircDiscoverer currently catalogs predicted CPI information for 445 unique proteins across all species analyzed by multistep strategies (See Materials and Methods), followed by summarizing 284 research articles (Figure 1). This enriched our database with a diverse and comprehensive set of CPIs and provided resources for experimental validation, as discussed below. Here, for each predicted circRNA, we defined two metrics: ***Sample_score***, which is the number of CLIP/RIP-seq samples of a given protein in which that circRNA is detected, divided by the total number of samples available for that protein, and ***circRNA_score,*** which represents the relative occurrence frequency of a circRNA within a CLIP/RIP-Seq dataset, and is calculated by dividing the number of samples in which a given circRNA is detected by the highest number of samples in which any circRNA is detected for that protein. A circRNA present in the maximum number of CLIP/RIP-Seq samples for a given protein gets the highest ***circRNA_score*** of 1, and the others were ranked relative to it. Both these scores normalized for differences in sample availability and avoided biases from raw counts due to cell-specific circRNA expression and cell-specificity CPIs(25, 26).

**Figure 1:**
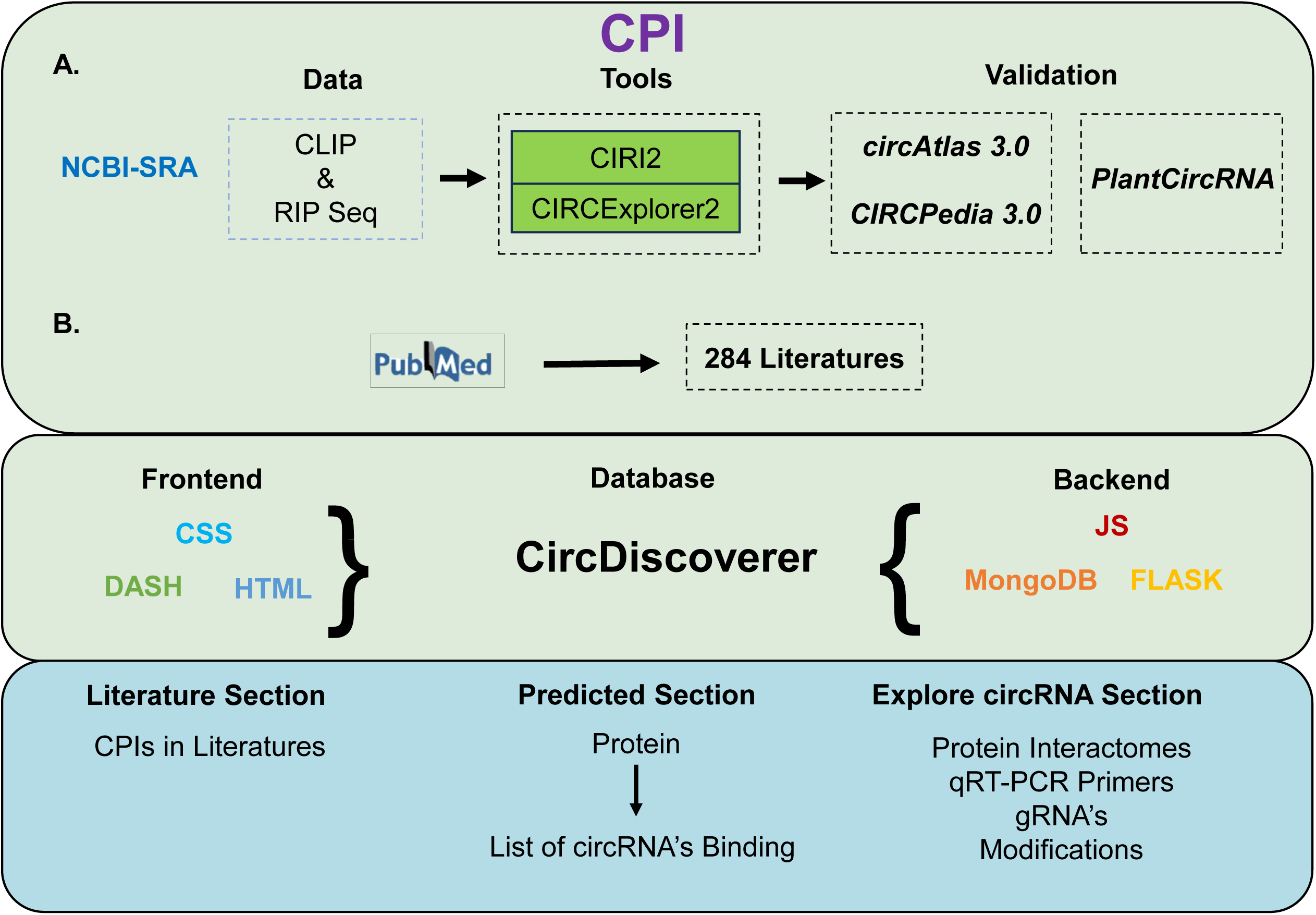
Workflow of the CircDiscoverer

### Validation-oriented functionality

A distinguishing feature of CircDiscoverer is its emphasis on experimental follow-up using integrated qRT-PCR primers and RfxCas13d gRNA design modules for validating CPIs. qRT-PCR primers can be used to validate CPIs through CLIP- and RIP-seq-based pull-downs(2, 27, 28). In parallel, RfxCas13d-compatible gRNAs can be incorporated into catalytically inactive CasRx fused with biotin ligase to target the back-splice junction (BSJ) of circRNAs for CPI investigation(19, 29). These gRNAs can also be used for circRNA knockdown to study the functional role of specific CPIs. This strategy is particularly useful because recent studies have shown that RfxCas13d-compatible gRNAs can achieve a higher knockdown efficiency than conventional siRNA- or shRNA-based approaches(22, 30).

Additionally, nine types of predicted RNA modifications in circRNAs, that is, m6A, m5C, m1A, ac4C, Pseudouridine, Nm, m7G, A-I, and modifications classified as other types, including m2G, m6Am, and m5U, are available along with the incorporation of experimentally validated RNA modifications from literature wherever available, thereby allowing CPIs to be linked with specific modification sites in circRNAs. Finally, CircDiscoverer records known protein-binding motifs for 59 proteins, allowing users to interpret CPI candidates in a mechanistic framework rather than as isolated associations.

### User interface and query modes

The web interface of CircDiscoverer consists of three main sections: Literature, Predicted, and Explore circRNA, which contains four more sections, as shown in Figure 2. We also have the Help page and download sections, containing detailed tutorial, contact information, and downloadable files. Shown briefly in the tutorial page with examples, (Additional file 2), in particular, the **<u>Literature section</u>** compiles experimentally validated CPIs from published studies across various diseases and physiological processes, while the **<u>Predicted section</u>** lists details of additional potential CPIs across the selected organism of interest, and the **<u>Explore circRNA section</u>** allows users to analyze a specific circRNA of an organism in more detail, such as its protein interactomes and gRNAs for cloning in RfxCas13d expressing plasmid vector called <u>pXR004</u>(31), qRT-primers, and the possible predicted and known experimentally validated modifications wherever available, since not all circRNAs possess modifications. In particular, the primers and guide RNA section lead users directly to the <u>NCBI-Primer BLAST</u> or <u>NCBI-BLAST</u> page, with the sequences autofilled, enabling users to verify off-target sequences. All circRNA data were mapped to their respective genome assemblies for standardization.

**Figure 2:**
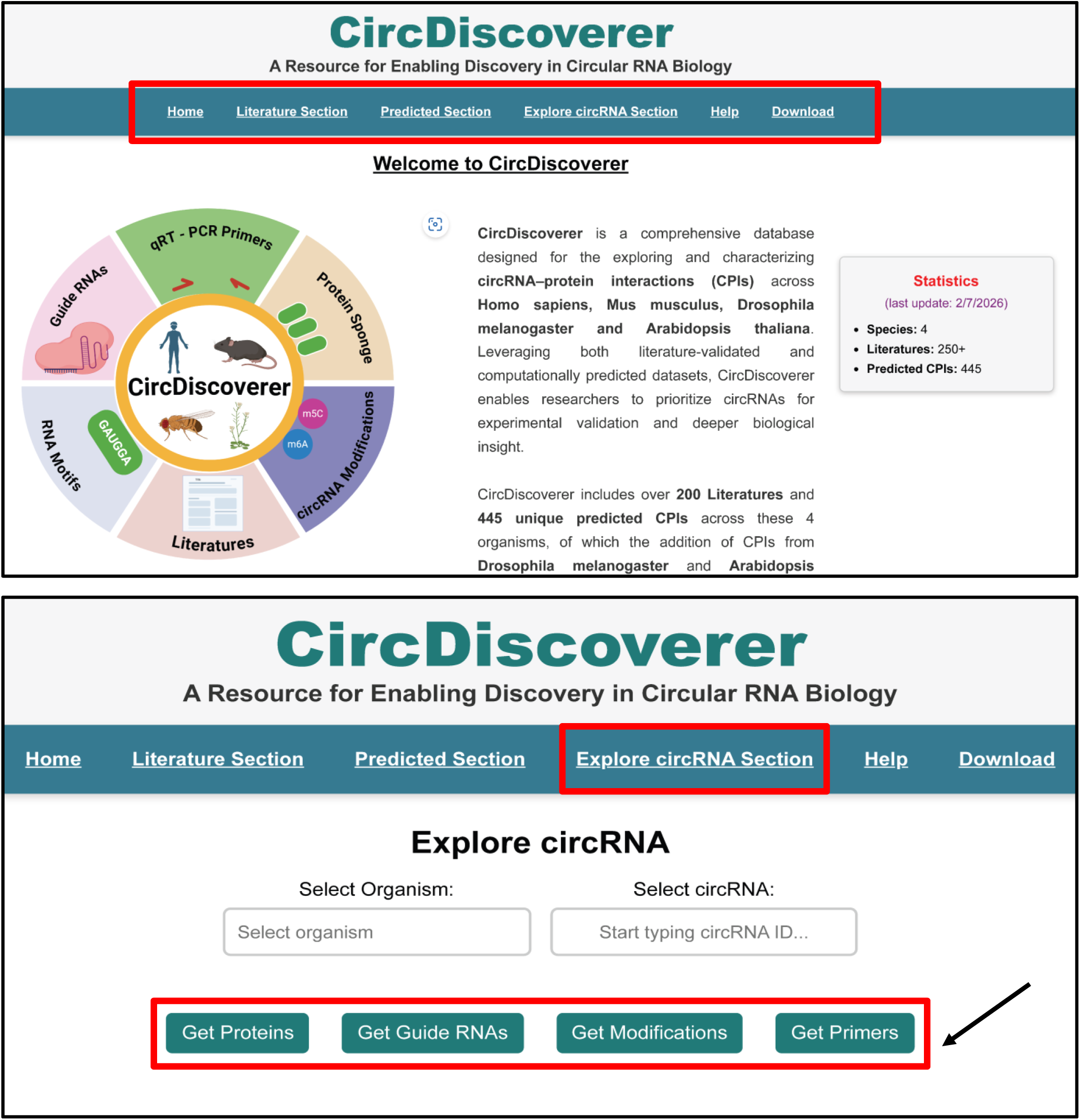
View of the CircDiscoverer website pages. A) The Home page and the sections inside it, B) The Explore circRNA sections and the tabs inside it

### Comparison with Other Databases

Of the several resources currently available for exploring circRNAs (Table 1), only a few of them cover CPIs and are often limited in the coverage of proteins (Table 2, Figure 3A) or in offering annotating resources to validate and verify these CPIs in a single platform (Figure 3C, 3D).

**Figure 3:**
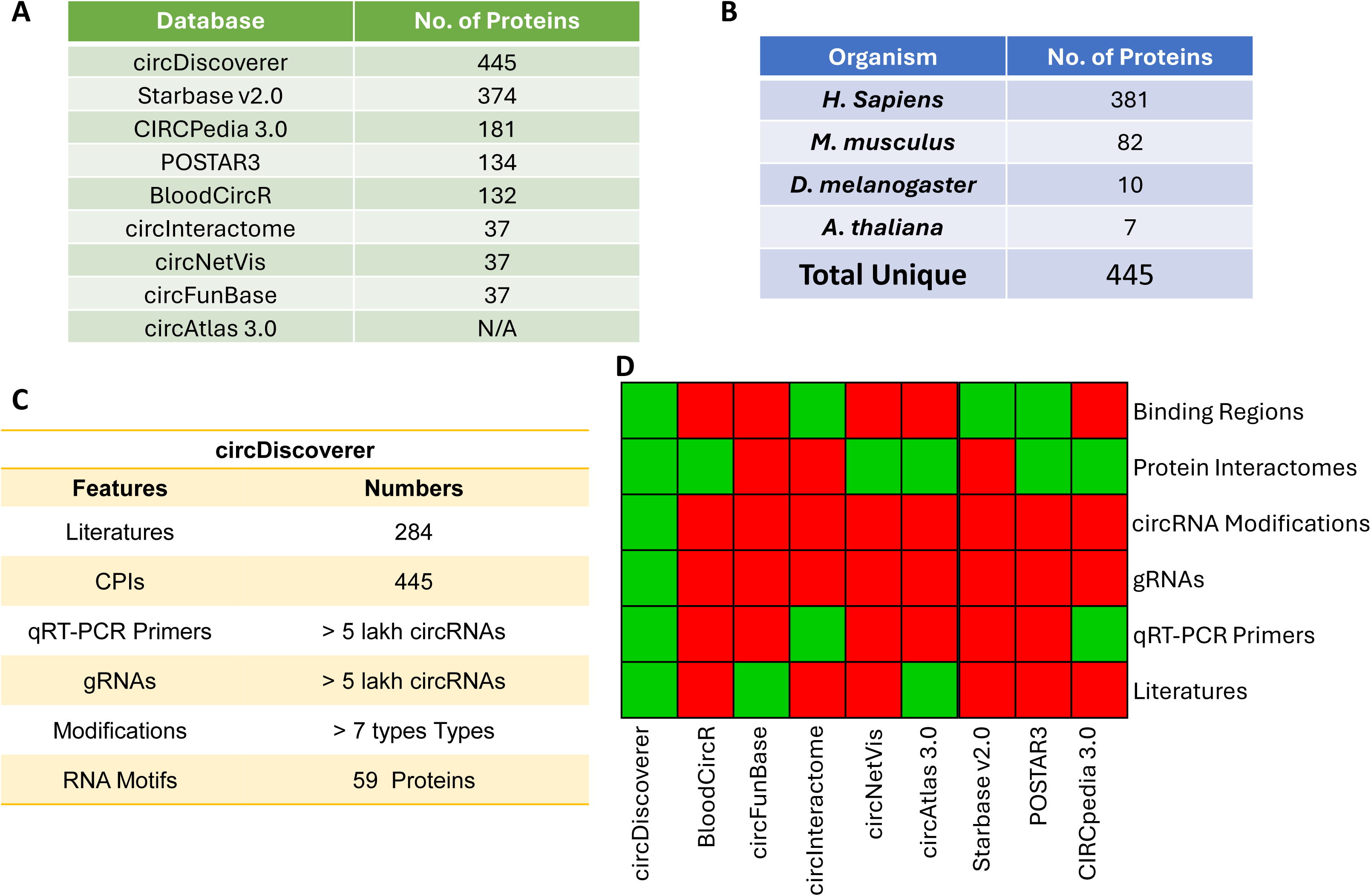
Statistics and Comparison of circDiscoverer with other databases. A) Comparison of the number of proteins in CircDiscoverer with other CPI centric databases, B) No. of organism specific proteins in CircDiscoverer, C) Features and Statistics of CircDiscoverer, D) Heatmap showing the absence and presence of an attribute in a database in Red and Green colors, respectively, thereby, specifying that, only CircDiscoverer holds all the attributes compared to other CPI-centric databases.

CircDiscoverer contains CPI information for 445 unique proteins across *Homo sapiens*, *Mus musculus*, *Drosophila melanogaster*, and *Arabidopsis thaliana* (Figure 3A, B). Compared with the largest currently available CPI resource, starBase v2.0 (J. H. Li et al. 2014), which contains CPI information for 272 human and 102 mouse proteins, CircDiscoverer not only covers proteins represented in existing CPI databases, but also includes 193 uniquely represented human proteins and 38 uniquely represented mouse proteins (Figure 4A and 4B). This breadth is notable because it covers proteins with well-characterized RBP’s(32), novel RBPs, and proteins yet not known to have any RNA-binding domain(33) by analyzing their CLIP-based evidence, RIP-based evidence, and motif-supported inference, rather than relying on a single evidence stream (Figure 5A, 5B).

**Figure 4:**
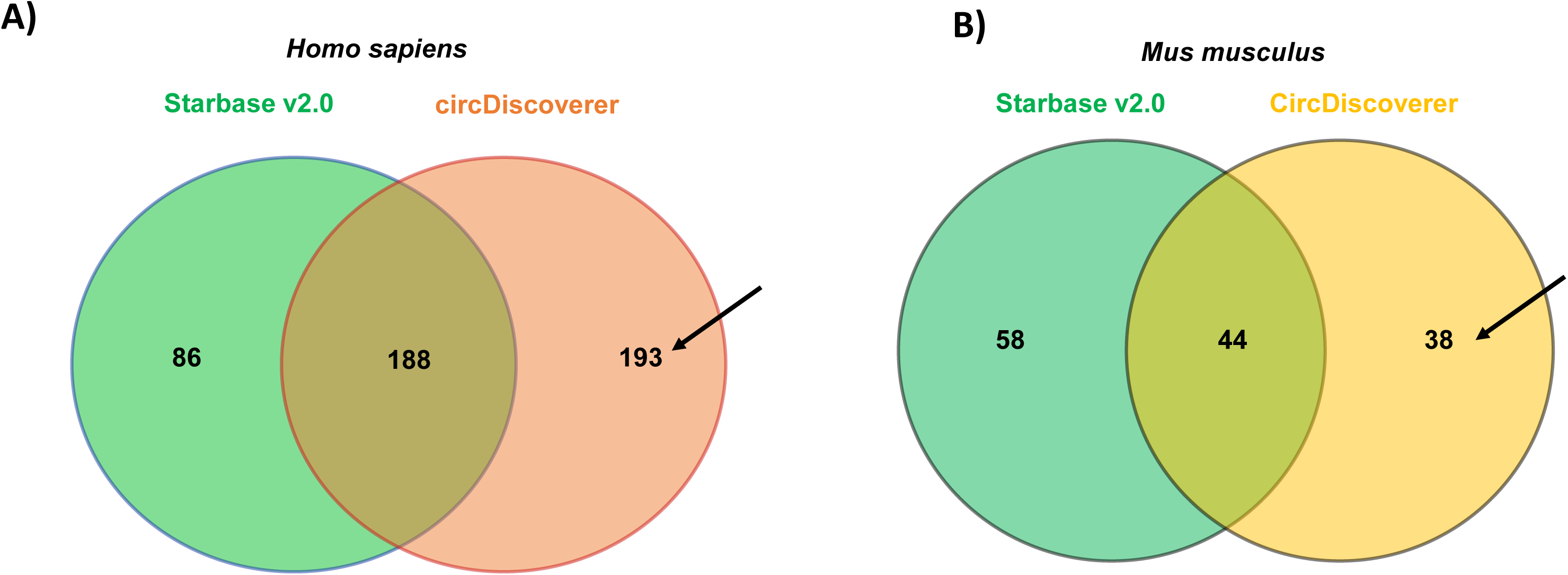
Comparison of No. of Proteins in CircDiscoverer with largest CPI Repository Starbase v2.0 for. A) Homo sapiens, showing 193 unique protein, B) Mus musculus, showing 38 unique proteins.

**Figure 5:**
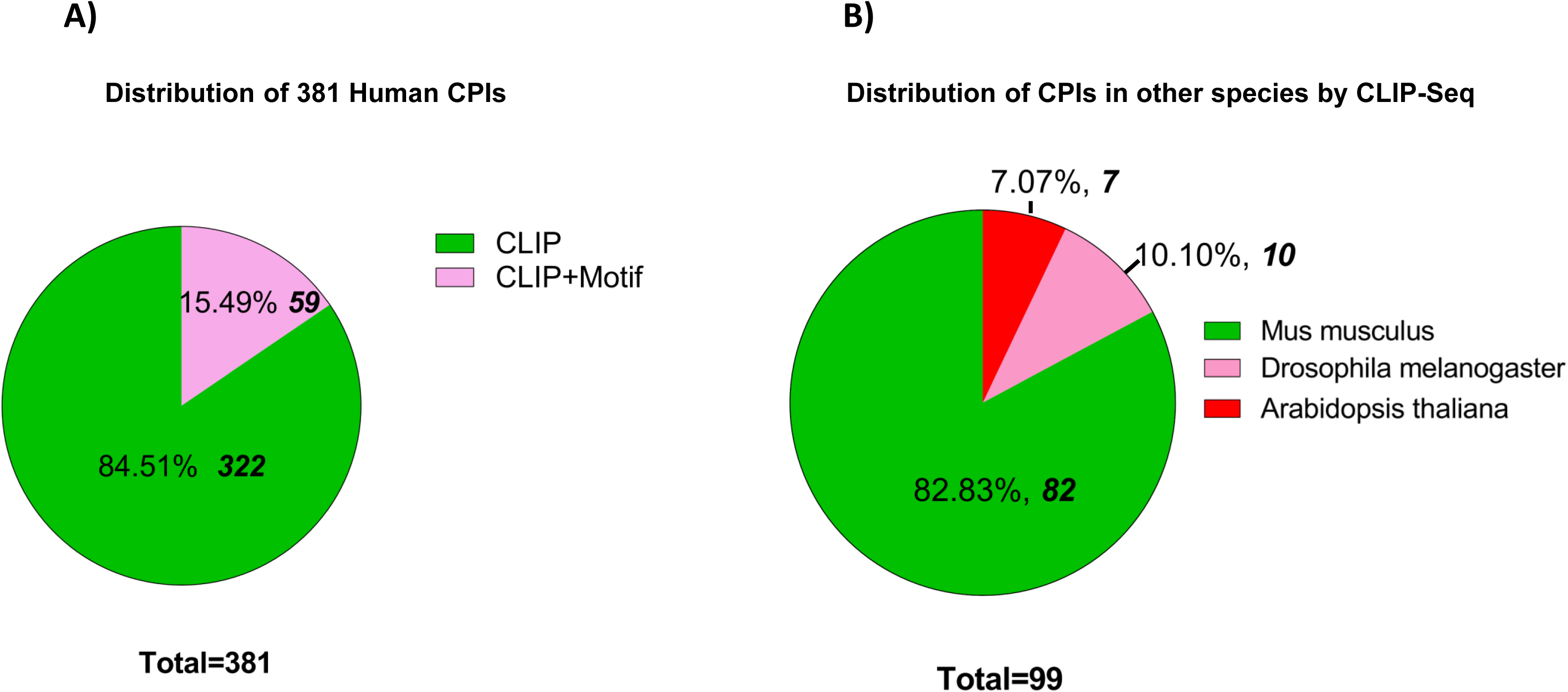
Source of Distribution of CPI in CircDiscoverer for. A) All the 381 CPI in *Homo sapiens*, B) For 99 CPIs across other species in CircDiscoverer by CLIP-Seq.

In addition to the predicted interactions, CircDiscoverer also summarized 284 articles of CPIs, offered annotation resources to validate and associate these CPIs in a single platform, and bridged a critical gap when compared to current resources (Figure 3C and 3D).

### Case Study

#### Case Study on CPI

SRSF1 is a well-characterized splicing factor that plays a crucial role in multiple biological processes and diseases. Particularly, in HIV-1, SRSF1 binds to the viral promoter and inhibits viral transcription(34). Querying SRSF1 in the Predicted Section of CircDiscoverer revealed several candidate circRNAs, of which we selected hsa_SMARCA5_0005, which had a good ***sample score***, and a good ***circRNA_Score***, as reported by two tools and two types of evidence, CLIP-Seq and motif-based evidence (Figure S1). Examining hsa_SMARCA5_0005 further in the Explore circRNA section showed that SRSF1 appeared as the top interacting partner, supported by a higher number of binding motifs (Figure S2). Cross-verification with the literature further confirmed that the hsa_SMARCA5_0005–SRSF1 interaction was experimentally validated(19), highlighting how CircDiscoverer enables users to link predictions with biological evidence and design experiments. Other case studies are presented in Table 3.

#### Case Study on circRNA modification

hsa-PUM1_0016 has been confirmed to influences non-small cell lung cancer (NSCLC) via an unknown mechanism(35). As CPI also depends on circRNA modification (36), we queried hsa-PUM1_0016, which predicted m6A modification (Figure S3A) and was also validated experimentally(35). Similarly, hsa-CCSER2_0002 was predicted to have m6A modification (Figure S3B) in CircDiscoverer, and it was experimentally shown to contain m5C modification(37). Since circRNA modification occurs in a cell-type-specific manner, it is possible that hsa-CCSER2_0002 might undergo m6A modification in other cell types(38), highlighting the possibility that proteins interacting with a circRNA can be associated with modifications inside the circRNA(32). All other case studies related to the modifications are listed in Table 4.

## 4. Discussion

Although circRNA biology has expanded rapidly, existing CPI resources remain limited in coverage and often lack sufficient annotations for experimental validation. CircDiscoverer expands this landscape by integrating literature-reported and computationally predicted CPIs with validation-oriented annotations, including qRT-PCR primer design, gRNAs, RNA modification information, and RNA-binding motifs. CircDiscoverer contains CPI information for 445 unique proteins across *Homo sapiens*, *Mus musculus*, *Drosophila melanogaster*, and *Arabidopsis thaliana*, substantially expanding the available CPI repertoire. In addition to species represented in major CPI repository (39), CircDiscoverer further incorporates extensive CPI-related information from *Drosophila melanogaster* and *Arabidopsis thaliana*, thereby extending CPI exploration beyond mammalian systems into invertebrate and plant models.

A major strength of CircDiscoverer is that it functions not only as a repository of interactions but also as an integrated platform for biological interpretation and experimental planning. This is particularly important, because one of the major challenges in CPI research is translating computational predictions into experimentally testable hypotheses. Beyond the predicted interactions, CircDiscoverer incorporates known literature evidence, curated and predicted qRT-PCR primers, guide RNAs, nine types of RNA modifications beyond m6A and m5C, and protein-binding motifs wherever available. For both gRNAs and qRT-PCR primers, users can directly evaluate potential off-targets through integrated links to NCBI Primer-BLAST and NCBI BLAST, with sequences automatically populated (Additional File 2). By integrating these features, CircDiscoverer supports hypothesis generation and downstream validation more effectively than databases limited to static interaction catalogs do.

CircDiscoverer is designed to support both experimental and computational research. For experimental biologists, the database provides qRT-PCR primers, guide RNAs, and streamlined off-target evaluations for CPI validation. For computational researchers, integrated CPI datasets provide a framework for network analysis, protein- and circRNA-centric prioritization, and future machine-learning approaches for CPI prediction and binding affinity estimation. The inclusion of multiple species further enables the comparative analyses of conserved and species-specific CPI networks across diverse biological systems.

Another strength of CircDiscoverer is the incorporation of context-rich annotations beyond the interaction data alone. RNA modification annotations are especially valuable because post-transcriptional modifications can influence RNA–protein interactions(32). Likewise, motif-level information for proteins with known RNA-binding preferences helps distinguish interactions supported by recognizable sequence features from those inferred primarily through functional genomic analyses. Together, these modules improve the utility of the platform for generating mechanistic hypotheses.

The case studies presented in this paper highlight these advantages. For example, the SRSF1–hsa_SMARCA5_0005(19) interaction demonstrates how CircDiscoverer can connect computationally prioritized candidates with existing biological evidence, thereby helping users identify biologically plausible and experimentally tractable interactions. Similarly, modification- and validation-focused examples show how users can examine circRNA modifications and directly retrieve primer or guide RNA designs for follow-up studies. This combination of interpretability and experimental accessibility strengthens the relevance of CircDiscoverer for studies on molecular mechanisms, disease biology, and biomarker discovery.

Despite these strengths, our study has several limitations. First, predicted CPI entries should be considered hypothesis generating rather than definitive evidence of binding. Second, both predicted and literature-reported CPI landscapes are likely to be cell-type- and condition-dependent, meaning that interactions identified in one context may not necessarily generalize across systems(26). Third, although the database provides extensive oligonucleotides, that is, primers and gRNAs, experimental validation is required. In some cases, owing to sequence constraints, non-standard rules were used to design oligonucleotides, which requires empirical optimization. Acknowledging these limitations, CircDiscoverer is a rigorous computational resource, rather than an overinterpreted interaction atlas.

Although CircDiscoverer incorporates multiple species, CPI datasets for species beyond *Homo sapiens* remain comparatively limited, owing to the restricted availability of publicly accessible data. We will continue to update and expand CircDiscoverer as additional datasets become available while also incorporating more organisms and virus-encoded circRNAs to enhance our resources. Furthermore, we will focus on integrating large language model (LLM)-based automated circRNA knowledge extraction to facilitate intelligent literature mining, structured information retrieval, and the dynamic summarization of emerging circRNA research.

CircDiscoverer provides a valuable resource for both computational and experimental RNA biology by integrating CPI discovery, literature curation, and validation-oriented annotation within a single platform. Its major contribution lies not only in expanding CPI coverage but also in linking interaction predictions with experimentally actionable information. Consequently, CircDiscoverer should facilitate studies focused on underexplored CPIs, conserved circRNA–protein regulatory mechanisms, disease-associated candidates, and experimentally testable circRNA–protein interaction hypotheses.

## Supporting information

Supplementary Figures S1 through S3

## Acknowledgements

This work was supported by the Indian Council of Medical Research [grant numbers CAR-2024-01-000113 to A.C., IIRPSG-2024-01-00869 to S.C. and A.C.], the Lady Tata Memorial Trust, the European Molecular Biology Organization, the Anusandhan National Research Foundation, the DBT-Wellcome Trust India Alliance, to A.C.

S.S. is supported by a fellowship from the Indian Institute of Science Education and Research Bhopal. S.K. is supported by a fellowship from the Translational Health Science and Technology Institute.

## Author contributions

Data curation: SS. Technical help and website design: SK and SS. Original manuscript writing: SS; manuscript review: AC, SK, and S.C. Conceptualization and supervision: AC and SC.

## Website Availability

CircDiscoverer is accessible at https://aclab.iiserb.ac.in.

## Supplementary Figure Legends

**Figure S1:** Screenshot of hsa-SMARCA5_0005-SRSF1 interactions, showing List of circRNAs binding to SRSF1,

**Figure S2:** Proteins binding to hsa-SMARCA5_0005.

**Figure S3:** circRNA Modifications for A) hsa-PUM1_0016 as input, B) hsa-CCSER2_0002 as input.

