## Supplementary Figures S1 through S3 for "CircDiscoverer: A multispecies comprehensive resource for circRNA–protein interactions and RNA modification landscapes"

Figure S1

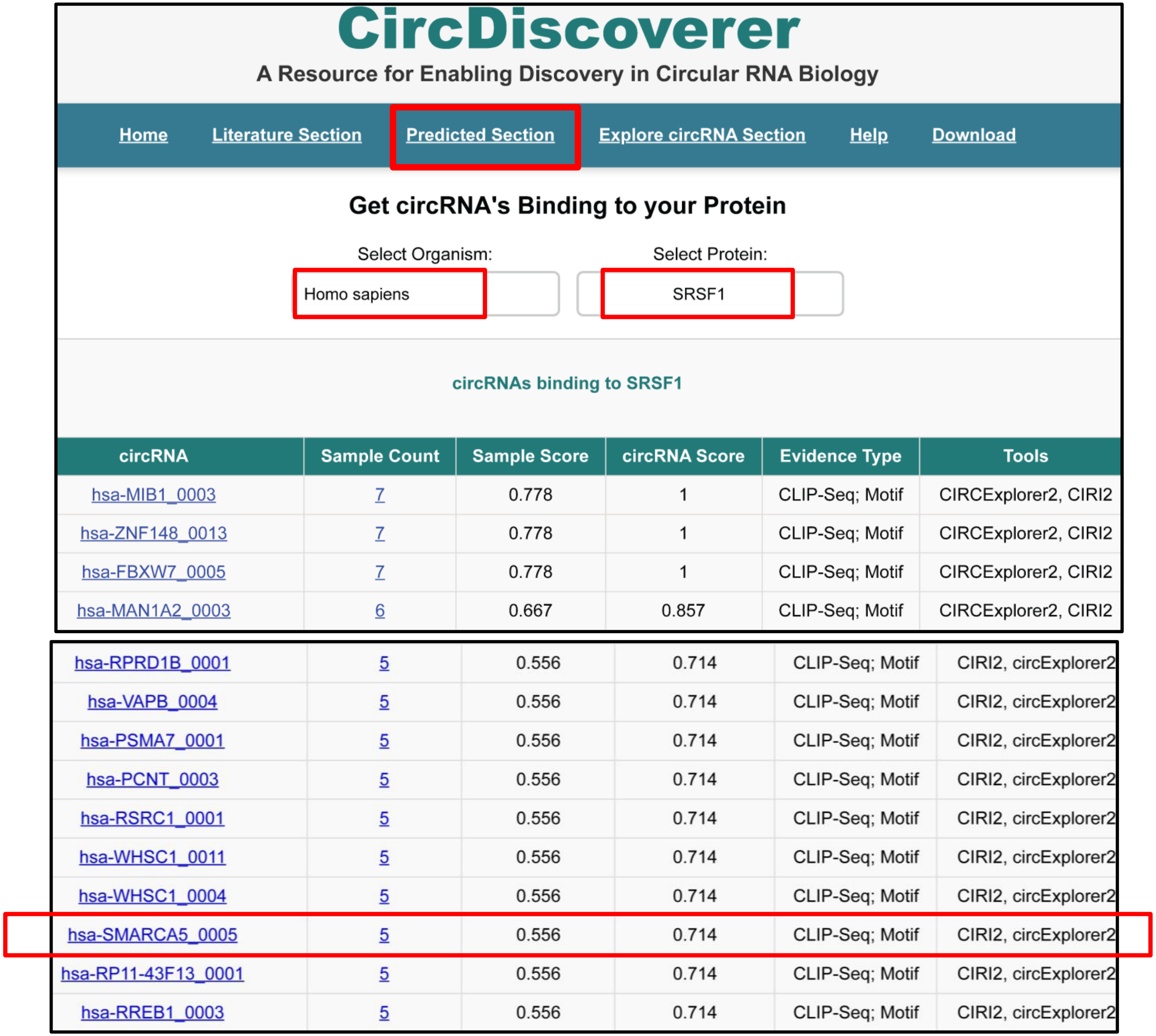

CircDiscoverer

A Resource for Enabling Discovery in Circular RNA Biology

Home | Literature Section | Predicted Section | Explore circRNA Section | Help | Download

Get circRNA's Binding to your Protein

Select Organism: Homo sapiens

Select Protein: SRSF1

circRNAs binding to SRSF1

| circRNA | Sample Count | Sample Score | circRNA Score | Evidence Type | Tools |
|---|---|---|---|---|---|
| hsa-MIB1_0003 | 7 | 0.778 | 1 | CLIP-Seq; Motif | CIRCExplorer2, CIRI2 |
| hsa-ZNF148_0013 | 7 | 0.778 | 1 | CLIP-Seq; Motif | CIRCExplorer2, CIRI2 |
| hsa-FBXW7_0005 | 7 | 0.778 | 1 | CLIP-Seq; Motif | CIRCExplorer2, CIRI2 |
| hsa-MAN1A2_0003 | 6 | 0.667 | 0.857 | CLIP-Seq; Motif | CIRCExplorer2, CIRI2 |
| hsa-RPRD1B_0001 | 5 | 0.556 | 0.714 | CLIP-Seq; Motif | CIRI2, circExplorer2 |
| hsa-VAPB_0004 | 5 | 0.556 | 0.714 | CLIP-Seq; Motif | CIRI2, circExplorer2 |
| hsa-PSMA7_0001 | 5 | 0.556 | 0.714 | CLIP-Seq; Motif | CIRI2, circExplorer2 |
| hsa-PCNT_0003 | 5 | 0.556 | 0.714 | CLIP-Seq; Motif | CIRI2, circExplorer2 |
| hsa-RSRC1_0001 | 5 | 0.556 | 0.714 | CLIP-Seq; Motif | CIRI2, circExplorer2 |
| hsa-WHSC1_0011 | 5 | 0.556 | 0.714 | CLIP-Seq; Motif | CIRI2, circExplorer2 |
| hsa-WHSC1_0004 | 5 | 0.556 | 0.714 | CLIP-Seq; Motif | CIRI2, circExplorer2 |
| hsa-SMARCA5_0005 | 5 | 0.556 | 0.714 | CLIP-Seq; Motif | CIRI2, circExplorer2 |
| hsa-RP11-43F13_0001 | 5 | 0.556 | 0.714 | CLIP-Seq; Motif | CIRI2, circExplorer2 |
| hsa-RREB1_0003 | 5 | 0.556 | 0.714 | CLIP-Seq; Motif | CIRI2, circExplorer2 |

**Figure S1:** Screenshot of hsa-SMARCA5_0005-SRSF1 interactions.

**Figure S2**

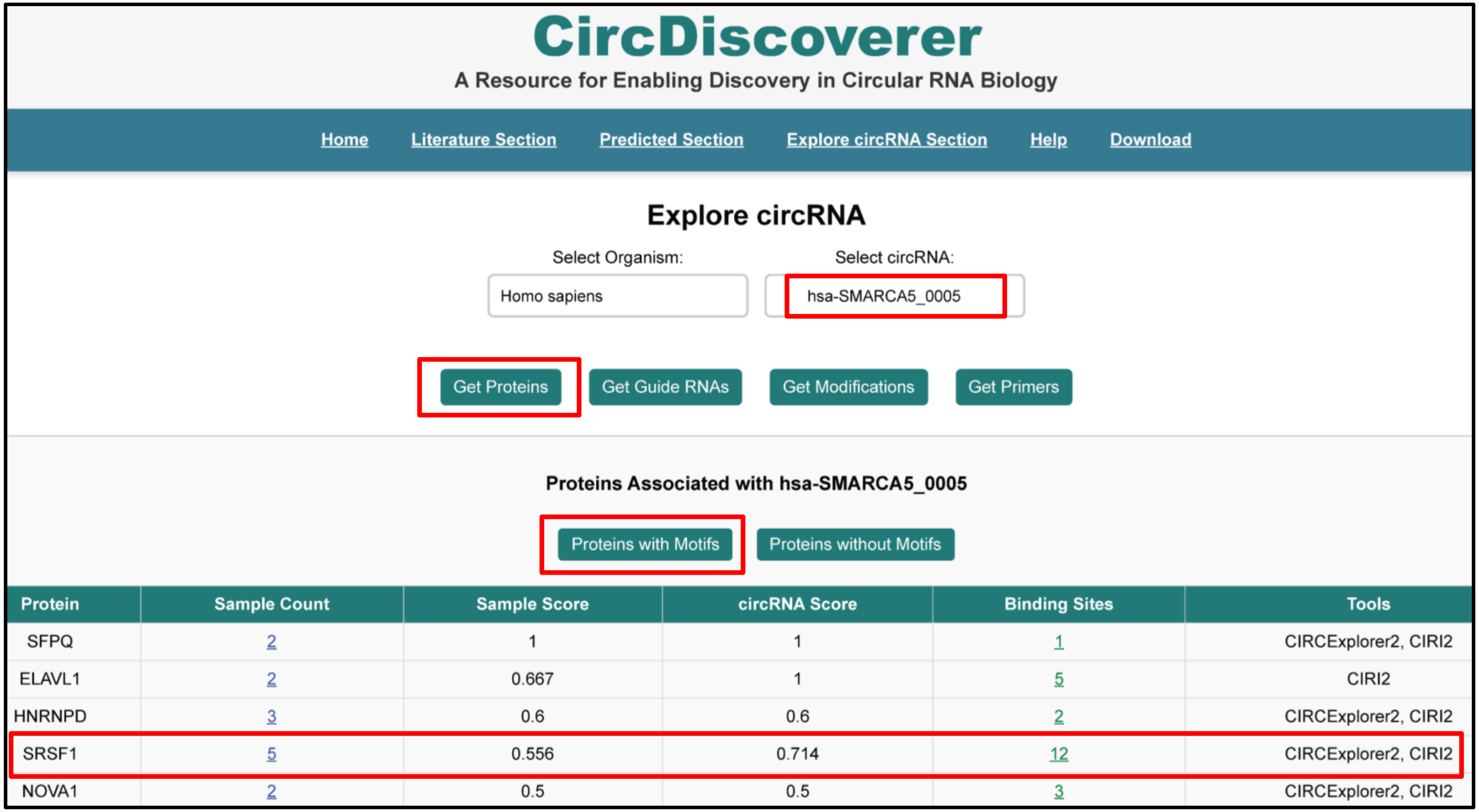


| Protein | Sample Count | Sample Score | circRNA Score | Binding Sites | Tools |
|---|---|---|---|---|---|
| SFPQ | 2 | 1 | 1 | 1 | CIRCExplorer2, CIRI2 |
| ELAVL1 | 2 | 0.667 | 1 | 5 | CIRI2 |
| HNRNPD | 3 | 0.6 | 0.6 | 2 | CIRCExplorer2, CIRI2 |
| SRSF1 | 5 | 0.556 | 0.714 | 12 | CIRCExplorer2, CIRI2 |
| NOVA1 | 2 | 0.5 | 0.5 | 3 | CIRCExplorer2, CIRI2 |

**Figure S2:** Screenshot of Proteins binding to hsa-SMARCA5_0005-with SRSF1 as one of top interactor

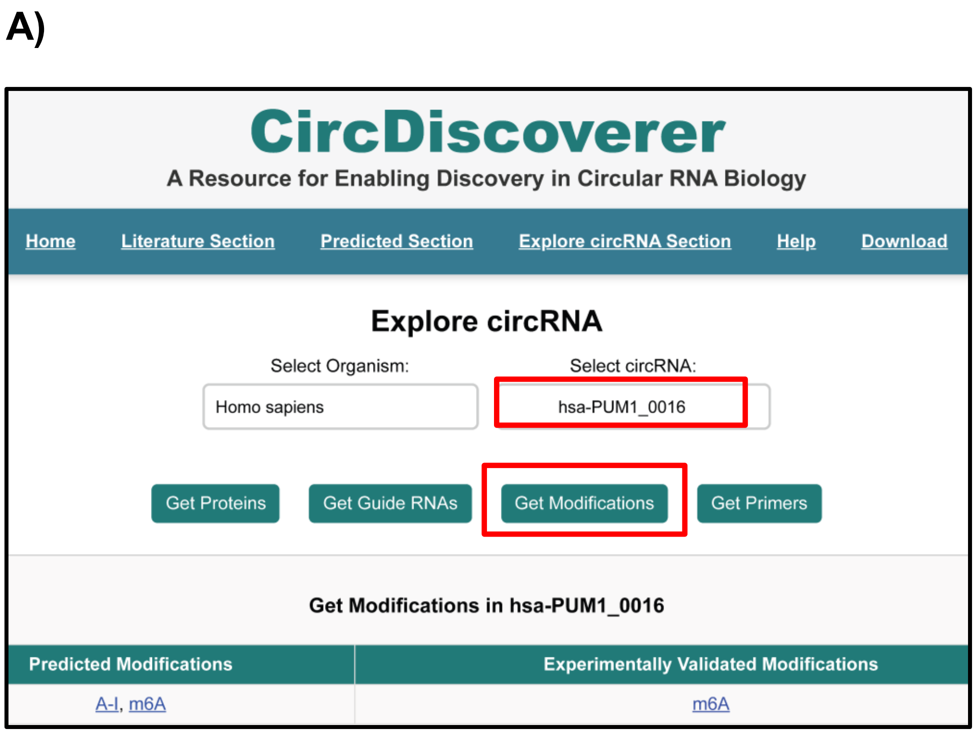


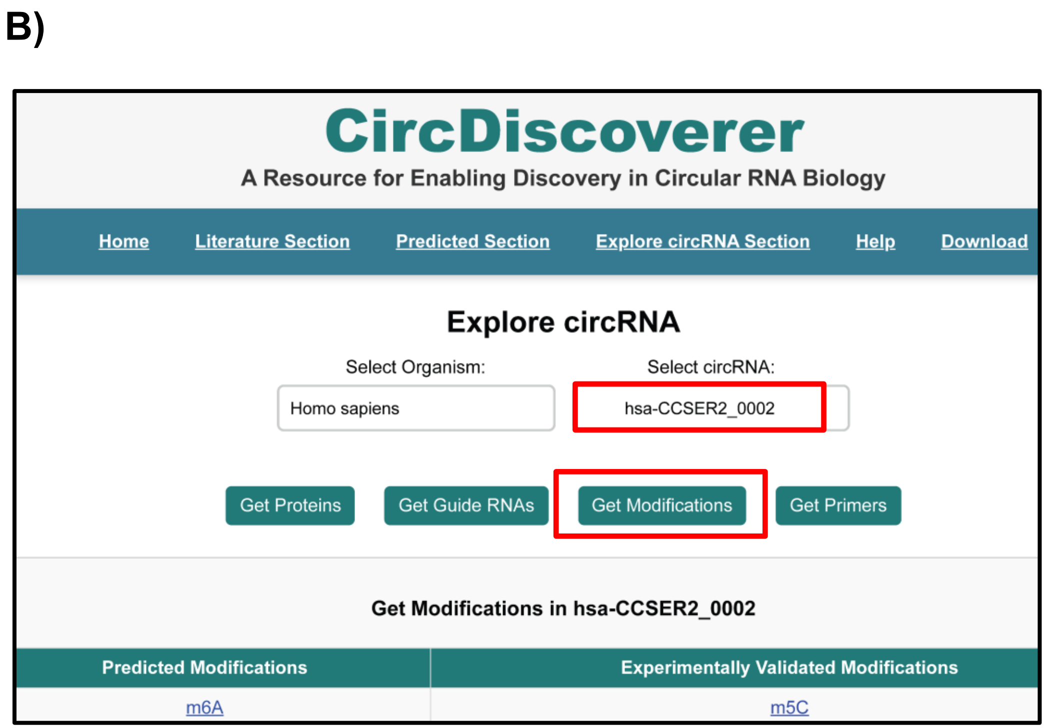


**Figure S3**: circRNA Modifications for A) hsa-CCSER2_0002 as input, B) hsa-PUM1_0016 as input.
